# Continuous dynamic modeling of regulated cell adhesion

**DOI:** 10.1101/582429

**Authors:** J. M. Ko, D. Lobo

## Abstract

Cell-cell adhesion is essential for tissue growth and multicellular pattern formation, and crucial for the cellular dynamics during embryogenesis and cancer progression. Understanding the dynamical gene regulation of cell adhesion molecules (CAMs) responsible for the emerging spatial tissue behaviors is a current challenge due to the complexity of these non-linear interactions and feedback loops at different levels of abstraction—from genetic regulation to whole-organism shape formation. Continuous mathematical models of cell adhesion are ideal for the modeling of the spatial dynamics of large cell populations, where different cell types define inherent adhesion strengths. However, biologically the adhesive properties of the cell arise dynamically from differential expression of CAMs, which are precisely regulated during development and cancer progression. To extend our understanding of cell and tissue behaviors due to the regulation of adhesion molecules, here we present a novel model for the spatial dynamics of cellular patterning, growth, and shape formation due to the differential expression of CAMs and their regulation. Capturing the dynamic interplay between genetic regulation, CAM expression, and differential cell adhesion, the proposed continuous model can recapitulate the complex and emergent spatial behaviors of cell populations that change their adhesion properties dynamically due to inter- and intracellular genetic regulation. This approach can demonstrate the mechanisms responsible for classical cell sorting behaviors, cell intercalation in proliferating populations, and the involution of germ layer cells induced by a diffusing morphogen during gastrulation. Integrating the emergent spatial tissue behaviors with the regulation of genes responsible for essential cellular properties such as adhesion will pave the way towards understanding the genetic regulation of large-scale complex patterns and shapes formation in developmental, regenerative, and cancer biology.

## 1. Introduction

The adhesive properties of cells can dictate their spatial behaviors and the formation of correct tissue patterns and shapes during morphogenesis and homeostasis (1). Seminal studies demonstrated how stirred disassociated embryonic tissues could sort themselves and regain their specific configurations due to the distinct adhesive properties of their different cell types (2, 3). These cell-cell adhesive forces are dependent on the expression of cell adhesion molecules (CAMs) through the cell surface, such as families of proteins including the cadherins, integrins, and nectins (4, 5). CAMs expressed at the cell surface can form bonds with the same or different CAM types expressed in neighbor cells, resulting in different adhesive strengths. These CAM adhesive forces are transmitted to the cell through its cytoskeleton network and can result in specific cell spatial behaviors. The sum of intercellular interactions between different CAMs determine the net force in the cell, which drive specific cellular movements and emergent tissue patterns. The importance of cell adhesion is clear when its cellular components are perturbed, resulting in tissues that can degenerate into mis-patterned phenotypes during development (6) and disease states such as cancer progression and metastasis (7, 8). However, the biophysical dynamics and cellular behaviors directed by differential adhesion and its genetic regulation are currently not completely understood.

The precise regulation of CAM expression modulates the adhesive properties of cells and hence can control the movement of cells and the formation of global tissue patterns during morphogenesis, whereas its dysregulation may lead to tumor formation and metastasis. Several gene families have been found to regulate CAM expression. The Snail family of transcription factors regulate the expression of cadherins essential for gastrulation in invertebrates, the epithelial-to-mesenchymal transition in neural crest cells in all amniotes, and the development of organs such as the kidney (9, 10). Differential regulation of CAMs such as cadherins by ephrins and *Hox* genes is a key factor for proper cell distribution during limb morphogenesis and regeneration (11); mutations in these pathways can result in limbs with abnormal morphological organizations (12). Dysregulated pathways controlling CAMs expression is sufficient to induce tumor progression, metastasis formation, and drug resistance (9, 13). Kinases can up-regulate E-selectin—a CAM essential for the localization of metastatic cancer cells in the lungs (14)—and specific kinase inhibitors targeting these pathways represent promising drugs for anticancer therapeutics (15). However, the complex feedback loops between CAM regulation, cellular adhesion dynamics, tissue behaviors, and intercellular signaling represents an extraordinary challenge that remains to be deciphered.

To understand the complex dynamics between the regulation of CAMs and the spatial tissue behaviors, mathematical and computational approaches are needed to model the physical properties of these processes and explain their emergent dynamics. Discrete models based on the extended Potts approach have been proposed to understand cell adhesion dynamics, and they can recapitulate the classical cell sorting dynamics due to adhesion (16–18), specific developmental dynamics (19–21), and cancer behaviors (22, 23). These models do not include the dynamics of CAMs expression, and instead use pre-defined adhesion constants for different cell types. Extensions to these discrete approaches have been proposed to model the concentration of CAMs, either using static concentrations defining cellular adhesion strengths (24) or dynamic concentrations with hybrid models (25). These approaches are based on the explicit modeling of cells, and hence computationally expensive for large numbers, which limits their applicability. In addition, mathematical methods to analyze discrete models are limited.

To overcome the limitations of discrete models, continuous models of cell adhesion have been proposed that can equally recapitulate the classical cell sorting behaviors but are computationally more efficient for the simulation of large populations and amenable for mathematical analysis (26, 27). Continuous models have been successfully used to explain developmental processes (28) and cancer dynamics (29–32). However, the adhesion properties in these models are static and defined with specific constants in pre-defined cell types. As a consequence, the regulation and dynamics of adhesion molecules have not been possible to model with continuous approaches, limiting our ability to understand the regulatory dynamics of CAMs expression and their influence in large scale tissue behaviors such as whole embryos.

Here we present a novel continuous model of cell adhesion due to the expression of CAMs and their regulation. This approach does not rely on pre-defined adhesion constants between cell types, but models as continuous the levels of CAM concentration, which in turn dynamically determine the adhesive properties between cells. Modeling the expression of CAMs naturally allows the inclusion of their regulatory dynamics, essential in many biological processes. We demonstrate the capabilities of the proposed model with three experiments. First, we show how the model can correctly recapitulate the classical Steinberg cell sorting dynamics due to the differential expression of CAMs. Next, we present a model of cellular intercalation dynamics resulting from the differential expression of two different nectins in a proliferating cell population. Finally, we model whole-embryo developmental dynamics during zebrafish gastrulation, showing how the diffusing morphogen Nodal regulates the expression of a cadherin, dynamically modulating the adhesive properties of cells and resulting in a characteristic involution of the mesendodermal germ layer. Integrating the regulatory dynamics of CAMs and their cell adhesion properties in the proposed continuous model permits the simulation and spatial predictions of the behaviors of large population of cells due to the interdependent dynamics of genetic regulation and adhesion proteins.

## 2. Model of dynamic cell adhesion

### 2.1. Model derivation

The dynamic adhesive properties of cells originate from the regulation and expression of CAMs. CAMs expressed on neighbor cells interact with each other, generating adhesive forces. CAMs bind to both CAMs of the same type, as well as CAMs of other types. The adhesive force generated from interactions between CAMs hence depends on both the adhesive strength between CAMs and their specific levels of expression in the interacting cells. The dynamic regulation of CAM expression, possibly by intra- and intercellular regulatory factors, results in dynamic adhesive forces. These dynamic forces can dictate cellular and tissue movement, resulting in target patterns and shapes. The proposed model follows a continuous approach to define a population of cells with adhesion forces (26, 27). The model does not explicitly define cell types, although it could, but instead the concept of cell type implicitly arises by the differential expression of factors and CAMs, which define the specific adhesive forces between cells.

We derive the model by considering the forces acting on a population of cells with no proliferation or death to be conservative, which implies by mass conservation

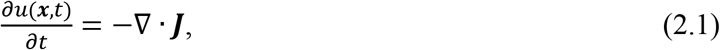

where *u*(***x***, *t*) is the cell density at position ***x*** and time *t*, and ***J*** is the flux of the cells. We can rewrite the cell density equation in terms of the flow velocity of the cells, resulting in

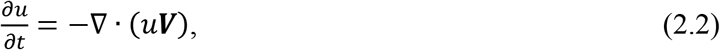

where ***V*** is the velocity field of the cells. Cells contain CAMs that are advected by the movement of the cells, resulting in

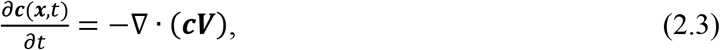

where ***c***(***x***, *t*) is the vector of CAM densities at position ***x*** and time *t*.

Cells move in a directed manner from regions of high density to those of lower density (27), causing dispersion velocity ***V***_***d***_, and towards each other due to adhesive forces between their expressed CAMs, causing adhesion velocity ***V***_***a***_, so the total velocity of the cells is

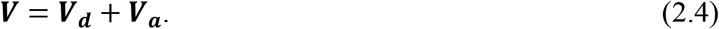

We assume that the cell dispersion velocity is proportional to the population density, which implies

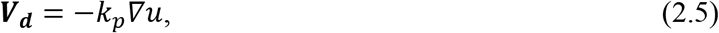

where *k*_*p*_ is the dispersion constant.

The adhesion velocity vector depends on the adhesive bonds between the CAMs expressed in the cells and their neighbors within a sensing radius *R* (Figure 1a). This radius models the size of a cell, including their ability to reach and contact other cells through the cell body and through their protrusions such as filopodia. Following Newton’s law and assuming that inertia is negligible for cell movements, the adhesion velocity vector is then inversely proportional to the cell radius (due to drag) and proportional to the sum of all adhesion forces between CAMs, such that

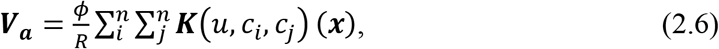

where *ϕ* is a constant of proportionality related to viscosity, *n* is the number of CAMs, and ***K***(*u*, *c*_*i*_, *c*_*j*_)(***x***) is the nonlocal cell adhesion force vector at location ***x*** due to the bonding interactions between CAMs *c*_*i*_ and *c*_*j*_. The adhesive strength between CAMs are defined by a symmetric square matrix *A*, where each element *a*_*ij*_ represents the adhesion strength between CAMs *c*_*i*_ and *c*_*j*_, and hence the diagonal defines the self-adhesion strengths for each CAM. The nonlocal cell adhesion force depends on the adhesive bindings between the CAMs expressed in the local cell at point ***x*** and those expressed by its neighbors within the cell sensing radius *R*. In *d* spatial dimensions, it takes the form

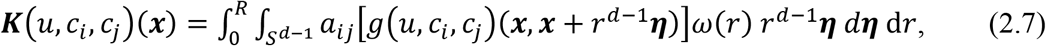

where *S*^*d*-l^ is the *d*-dimensional unit spherical surface, *r* is the radial distance, ***η*** is the direction vector, *g*(*u*, *c*_*i*_, *c*_*j*_)(***x***, ***x*** + *r*^*d*-l^***η***) describes the nature of adhesive forces between CAMs *c*_*i*_ and *c*_*j*_ expressed from cell locations ***x*** and ***x*** + *r*^*d*-l^***η***, respectively, and their dependence on the cell density, and *ω*(*r*) describes how the cell adhesive force depends on the radial distance *r*. For simplicity, we assume *ω*(*r*) = 1 in this paper.

**Figure 1.**
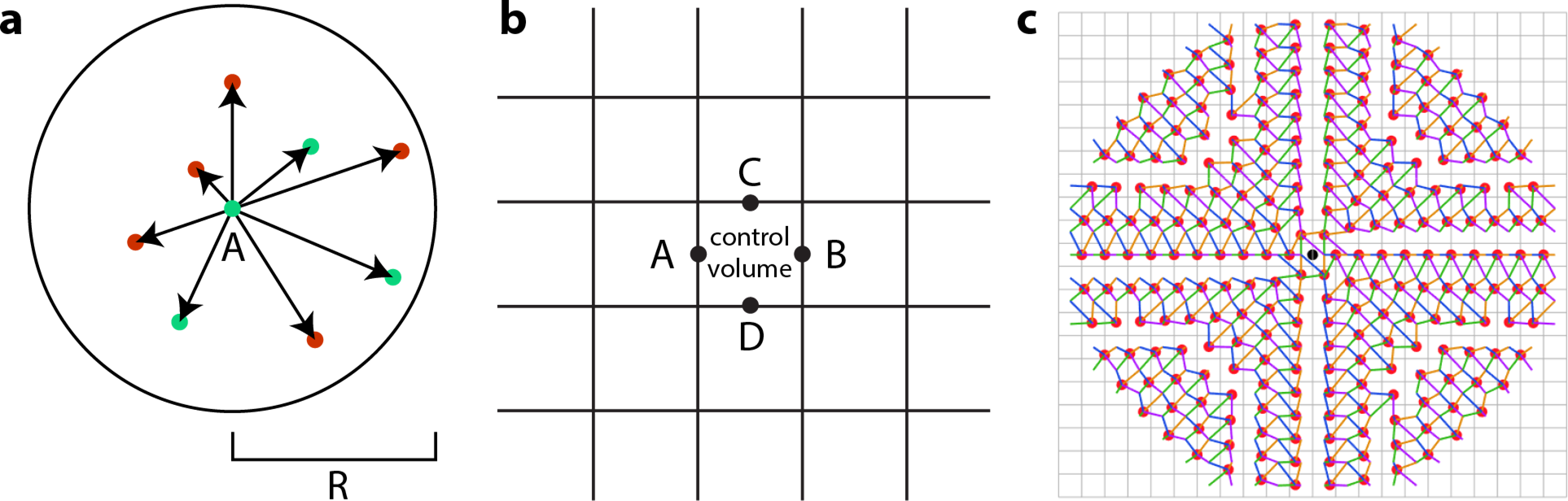
Proposed continuous model for the regulatory dynamics of cell-cell adhesion. **(a)** Cells regulate and express different adhesion proteins (CAMs, red and green), causing cell-cell adhesive forces depending on the CAMs concentration in the cells within a radius R. **(b)** Two-dimensional scheme of tissue discretization and cellular fluxes due to dynamic adhesion. Cell density and CAMs and other factors concentrations are defined in a grid of discretized control volumes, while the flux is defined across the boundaries between control volumes (points A-D). **(c)** Kernel for the numerical discretization in two dimensions of the adhesion integral at boundary point A in (a). The adhesion values are computed at points at regular angular and radial directions (red circles) from the boundary point (black circle at the center) with a bilinear interpolation of CAM concentrations from the center of the four surrounding control volumes (cyan, magenta, yellow, and green lines). The same kernel is used for boundary point B, while its transpose is used for boundary points C and D. The example shows a discretization with 42 angular by 10 radial directions.

The adhesive force between two CAMs expressed by two cells depends on their binding activity due to the CAMs relative concentrations within each cell. We assume that the binding activity follows the law of mass action, such as the adhesive force exerted on cells at location ***x*** expressing CAM *c*_*i*_ by cells at location ***y*** expressing CAM *c*_*j*_ depends on the product of their concentrations within their respective cells, given by

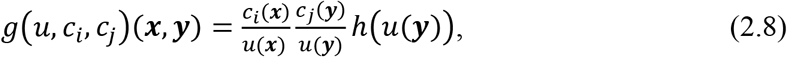

where *h*(*u*(***y***)) represents how the adhesive force depends on the local cell density. We assume a crowding capacity *k*_*c*_ in the population limiting the cell movement due to adhesion towards dense regions, such as

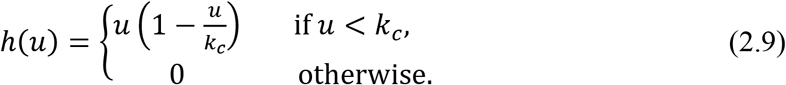

A nondimensional model is defined by rescaling with

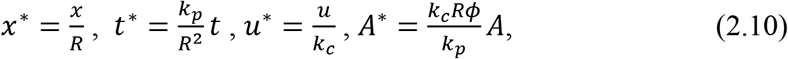

and dropping the stars, we obtain the model

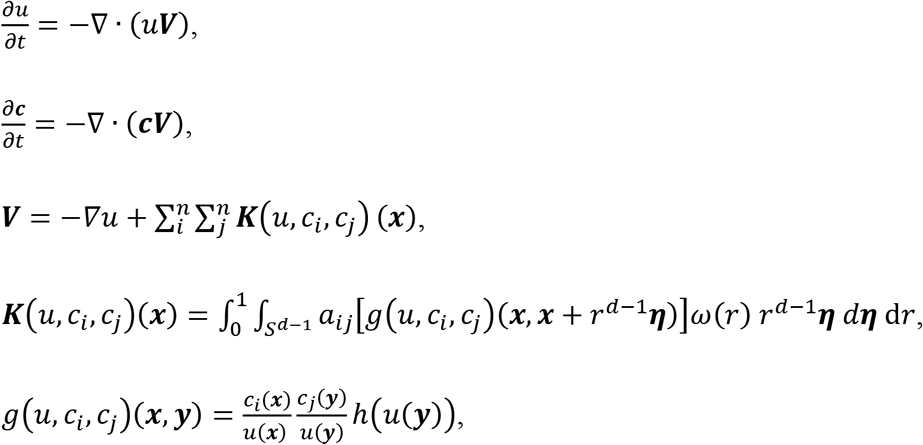

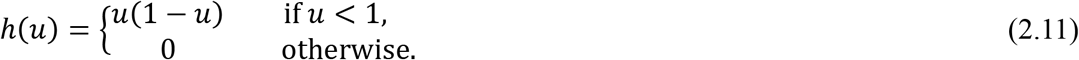

### 2.2. Numerical methods

PDE simulations were performed in a two-dimensional domain using the explicit upwind finite volume method with flux limiting in a uniform square lattice and a zero flux boundary condition. The fluxes between control volumes due to the adhesion velocity (2.6) are computed at four points *A* – *D*, one at each face midpoint (Figure 1b). At each of these midpoints and for each pair of CAMs, the nonlocal integral term for adhesion (2.7) is computed within a circle with radius *R* and centered at the face point (Figure 1c). The integral circle is discretized with parameters *N*_*r*_, *N*_*θ*_ ∈ ℕ, defining a set of points uniformly distributed along *N*_*r*_ radial values and 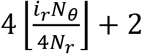 angular values for each radial value *i*_*r*_ ∈ ℕ, 1 ≤ *i*_*r*_ ≤ *N*_*r*_, as in (27). Since the cell density and CAM concentrations are numerically defined at the control volume centers, bilinear interpolation is used from the four surrounding control volume centers to calculate their values at the regular integral points in the circle (red points and color lines in Figure 1c) and the average from the two surrounding control volume centers to calculate their values at the face point (black point in Figure 1c). The bilinear interpolations of the integral circle are precomputed in a weight matrix representing a kernel, which then are used to efficiently calculate the adhesion velocities in each face midpoint with a kernel convolution operation (due to symmetry, the kernel for points *C* and *D* is the transpose of the kernel for points *A* and *B*). Cell densities and CAM concentrations outside of the domain are considered zero in the kernel convolution operation, as is consistent with the zero flux boundary condition (33). The system was numerically solved with a generalized Runge-Kutta fourth-order solver using ROWMAP (34). Simulation computations used MATLAB R2017b (The MathWorks, Inc.).

## 3. Simulations

We demonstrate the ability of the proposed model to recapitulate tissue shape behaviors due to the differential expression of CAMs with three simulations of *in vitro* and *in vivo* experiments. Classical cell sorting behaviors can be recapitulated in the model in a population of cells expressing two different CAMs, resulting in engulfment, mixing, or sorted cellular aggregates depending on the adhesive strengths between the expressed CAMs. Extending the model with cell growth, a simulation of *in vitro* growing dynamics shows how a proliferating cell population can result in either separated or intercalated patterns due to the cells expressing either the same or different nectins, respectively. Finally, the dynamics of zebrafish gastrulation are explained with an extended model including the expression of a morphogen forming a gradient, which in turn up regulates the expression of cadherins inducing the involution of these cells due to their acquired differential cell adhesion properties. Importantly, the behaviors shown in the simulations are not due to inherent cell adhesion strengths between different cell types, but from dynamic adhesion strengths that arise from the concentration of various CAMs, where each CAM type has molecular adhesion values and their concentrations can be subject to genetic regulation.

### 3.1. Cell sorting behaviors

CAMs can bind to each other with different adhesive strengths, so cell-cell adhesion forces depend on the levels of expression of the different CAMs. These differences in cell-cell adhesion can cause an emerging cellular self-organization into different spatial patterns, a behavior demonstrated *in vitro* in a variety of animal cells, including amphibian (2), chick (35), zebrafish (18), and hydra (36). Figure 2 shows how *in vitro* stirred suspensions of disassociated neural and epithelial retinal cells from a chick embryo can self-sort, forming two engulfed aggregates (37). The two cell types express different CAMs with different adhesive properties: CAMs in retinal epithelial cells self-adhere more strongly than CAMs in retinal neural cells, while the inter-adhesion between the two CAMs has an intermediate strength. The differential expression of these CAMs causes the cells to form aggregates, and since the CAMs expressed by the retinal epithelial cells adhere more strongly, these cells form tightly adhered aggregates, which hence are surrounded by the less strongly adhered retinal neural cells.

**Figure 2.**
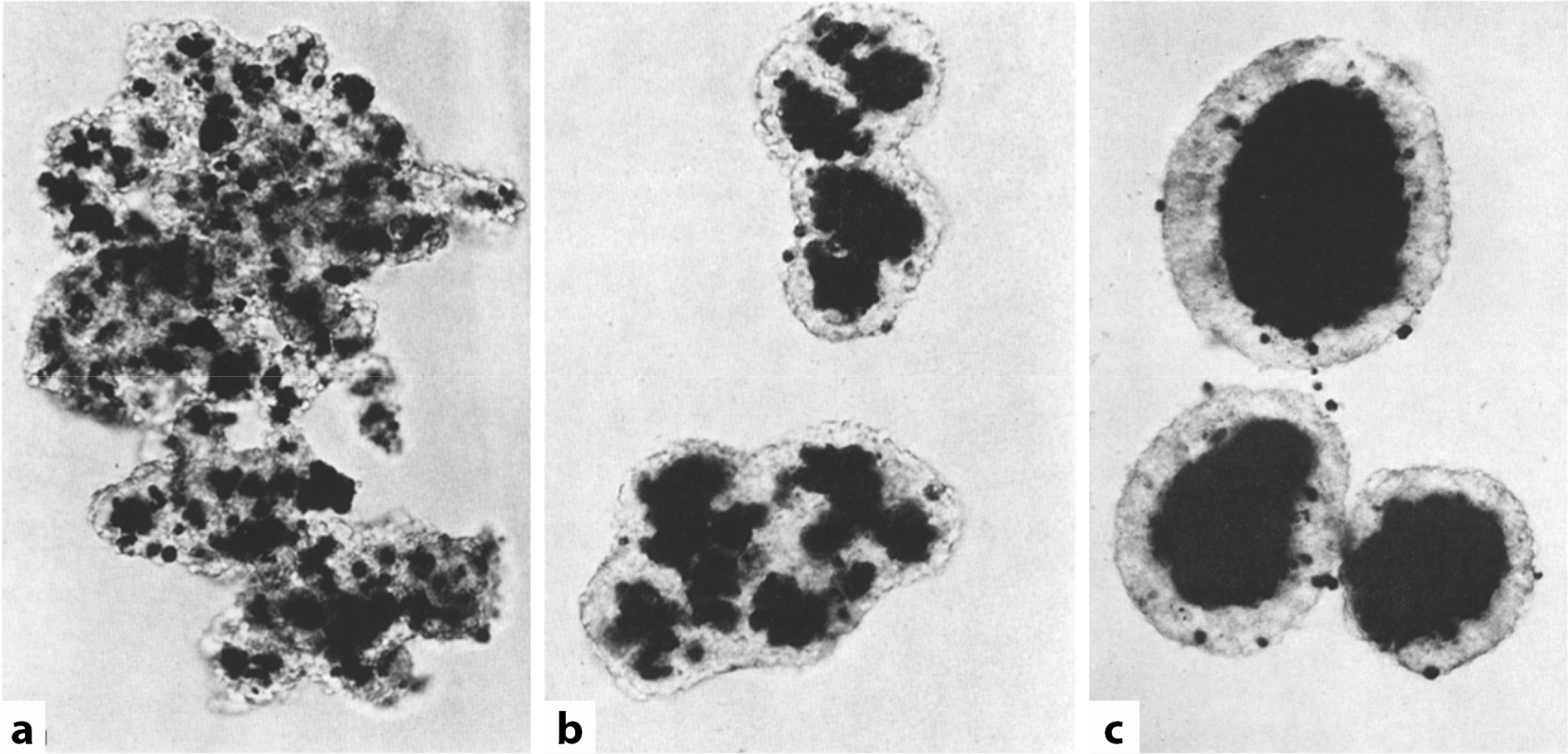
Cell sorting behavior in stirred suspensions of neural retinal cells (colorless) and pigmented retinal epithelial cells (black) from a seven-day old chick embryo. **(a)** Disordered mix after 5 h. **(b)** After 19 h., retinal cells form irregular shapes surrounded by neural cells. **(c)** After two days, retinal epithelial cells form round aggregates surrounded by retinal neural cells. Adapted from (37).

The proposed continuous nondimensional model (2.11) can recapitulate these cellular sorting behaviors due to the differential expression of CAMs. Figure 3 shows four different sorting behaviors resulting from non-proliferating cells expressing either of two CAMs with different adhesive strengths (strength values as in (27)). All the simulations start with the same initial random configuration of disassociated tissue, where each initial aggregate contains cells expressing one of two different CAMs. Depending on the relative strength of self- and cross-adhesion forces between the CAMs, the spatially-randomized tissues form aggregates that self-organize into patterns of engulfment, partial engulfment, mixing, or complete sorting. When the CAM self-adhesive strengths are asymmetric, similar to the retinal epithelial and retinal neural cells (Figure 2), the simulation recapitulates the engulfment behaviors observed *in vitro* (Figure 3a). This sorting behavior is due to the differential expression of CAMs, where cellular aggregates expressing the CAM with stronger self-adhesive force (red) are tightly adhered, and hence surrounded by the cellular aggregates expressing the CAM with weaker self-adhesive force (green). However, when the self-adhesive strength of the two CAMs are equal, but still higher than the cross-adhesive strength, no cell aggregate is stronger than the other, and hence there is still sorting between the tissues expressing the different CAMs, but no engulfment (Figure 3b). In contrast, the randomized tissues do not sort themselves when the cross- and self-adhesive strengths are equal (Figure 3c), resulting in aggregates that are mixed. In the completely absence of cross-adhesive strength between the two CAMs, the tissues expressing the different CAMs sort themselves completely, forming separated aggregates (Figure 3d). These simulations show how the cell sorting behaviors depend on the self- and cross-adhesion strengths between the CAMs and their levels of expression in the different tissues.

**Figure 3.**
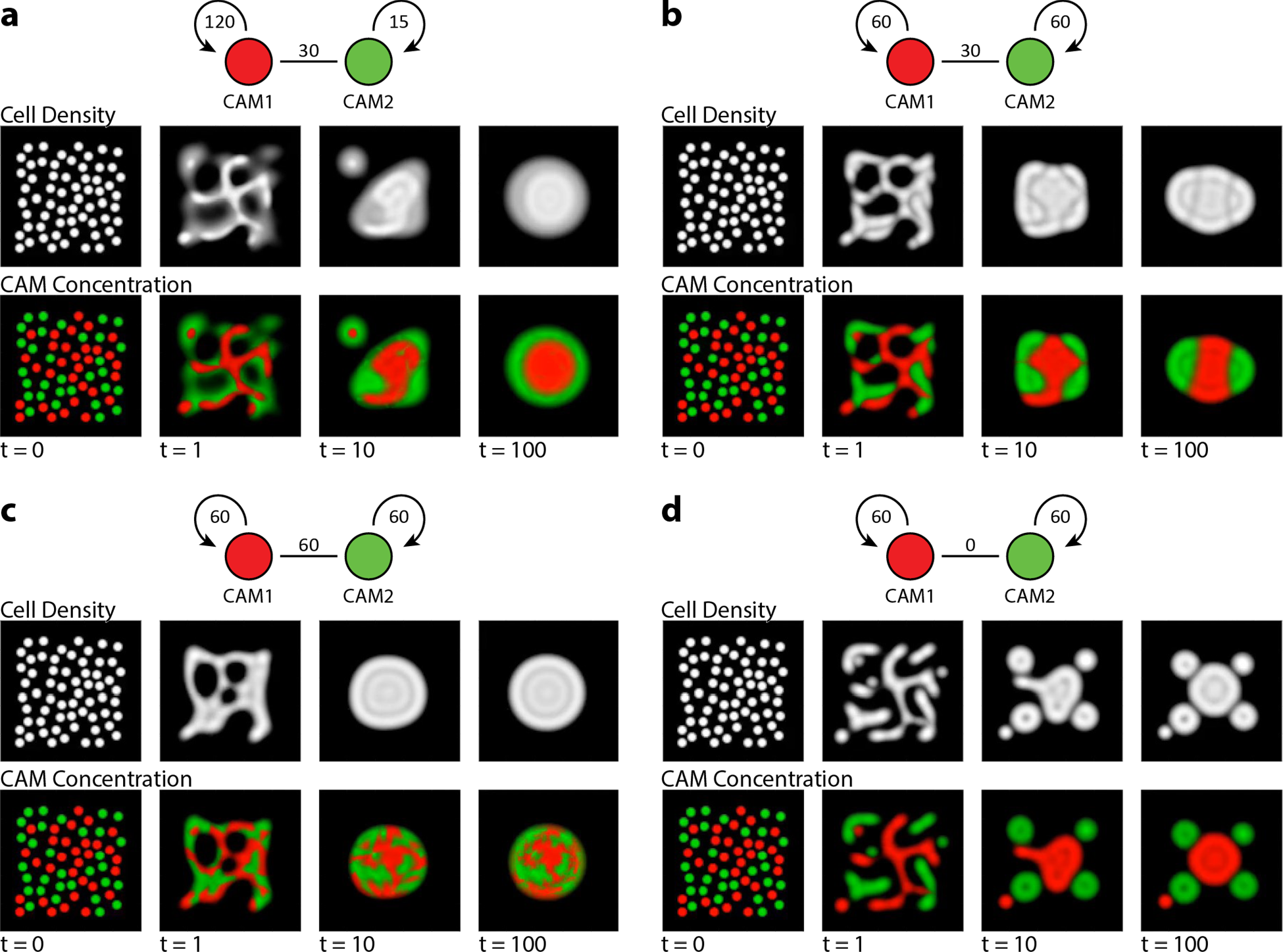
Cell sorting simulations in a population of cells expressing either of two CAM proteins with different adhesion strengths. **(a)** Asymmetrical self-adhesion strengths in the CAM proteins result in the engulfment of the cells expressing the higher adhesive strength protein (red) by the cells expressing the lower adhesive strength protein (green). **(b)** Symmetric protein self-adhesion strengths higher than the cross-adhesion strength result in partial engulfment. **(c)** When the protein self-adhesion and cross-adhesion strengths are equal, the cells become mixed. **(d)**Without cross-adhesion forces, the cell population expressing the two different CAM proteins completely sort themselves. Graph diagrams indicate protein self- and cross-adhesion strengths. All simulations start with the same initial state of equal red and green CAMs total concentration. Domain of size 10 × 10 units, discretized into a 100 × 100 grid. Arbitrary units.

### 3.2. Cell intercalation in proliferating cell populations

The spatial tissue behaviors in a population of proliferating cells can depend on both the expression levels and the adhesive properties of CAMs. This has been shown in *in vitro* assays of proliferating cell populations expressing similar or different nectin adhesion proteins (6). Figure 4 shows a co-culture assay of two separated growing populations of human embryonic kidney cells, each population transfected with similar or different CAMs in addition to two different fluorescent markers. When both populations express nectin-1, the boundary formed between the two populations at the contact plane is well defined, and the cells do not mix (Figure 4a). In contrast, when each population express a different CAM with different adhesive properties (nectin-1 or nectin-3), the two population mix and intercalate at the boundary (Figure 4b). The proposed model can explain these behaviors due to the differential expression of nectin-1 or nectin-3 in the two proliferating cell populations.

**Figure 4.**
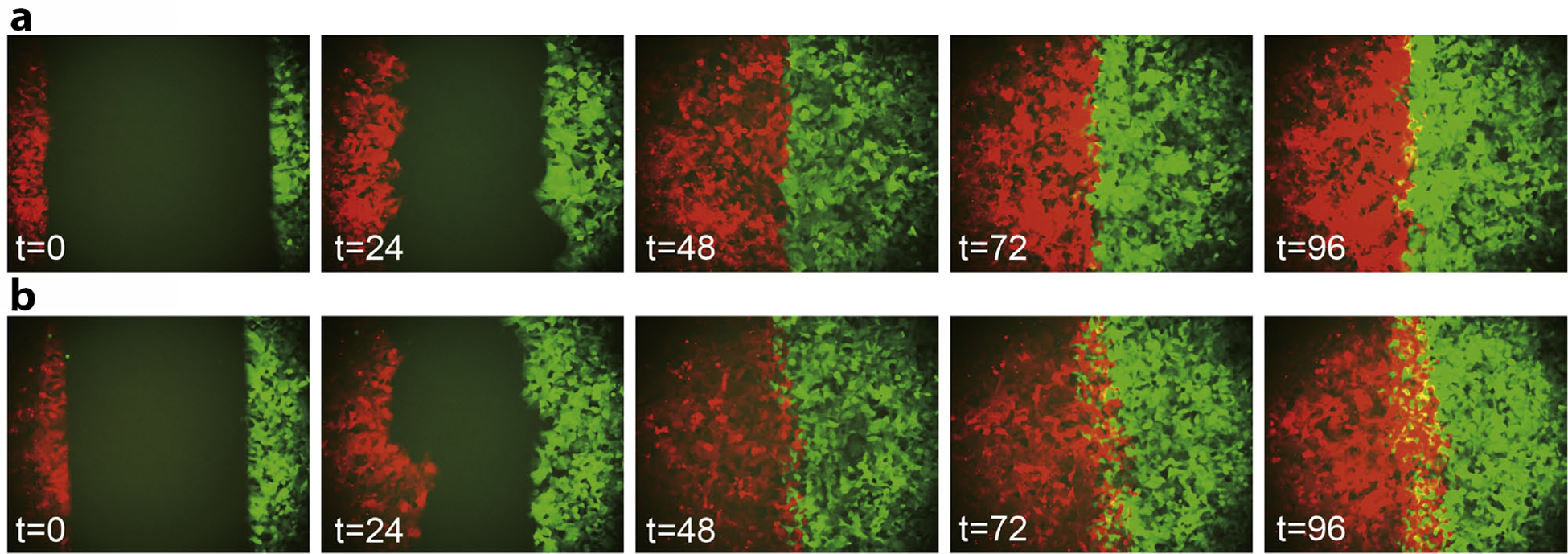
*In vitro* intercalation assay of co-culture of proliferating human embryonic kidney cells (HEK293) expressing similar or different CAMs, in addition to markers mCherry (red) or EGFP (green). **(a)** Cell populations expressing the same CAM, nectin-1 (red and green), do not mix at the boundary. **(b)** Cell populations expressing different CAMs, nectin-1 (red) and nectin-3 (green), mix and intercalate at the boundary. Time in hours. Adapted from (27).

We extend equations (2.2) to include a simple logistic cell growth term *g*(*u*), such that

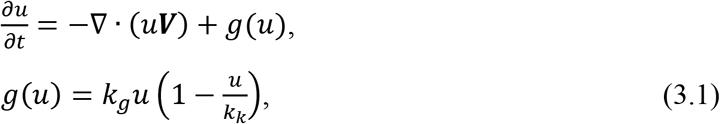

where *k*_*g*_ is the cell growth rate and *k*_*k*_ is the cell carrying capacity. We assume that the daughter cells continue expressing the same CAMs than their parent cells, so that the relative CAM concentration in the daughter and parent cells are equal. In this way, we extend equation (2.3) to

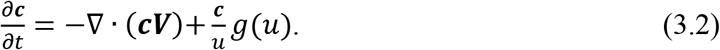

This extended model can recapitulate the *in vitro* intercalation dynamics of growing cell populations resulting from the differential expression of CAMs. We use the dimensional parameters as experimentally measured in (27):

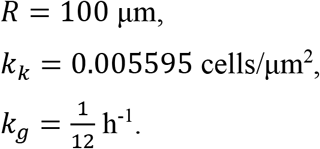

The parameter *k*_*c*_ could not be measured experimentally and was set to *k*_*k*_. We then fit the model to the experimental images from (27), setting *ϕ* = 800 and *k*_*p*_ = 200 μm^2^/h · μm^2^/cells.

Figure 5 shows simulations of growing population dynamics when the two cell populations express either the same (nectin-1) or different (nectin-1 or nectin-3) CAMs. The self- and cross-adhesion strengths of nectin-1 and nectin-3 are derived from protein-protein interaction experimental data measured with surface plasmon resonance (38) and their values are shown in Figure 5a. Both simulations start with the same random cell density (to avoid unstable steady states) and homogeneous relative CAM concentration 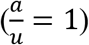. During the simulation, the cells proliferate and spread through all the domain up to the carrying capacity density. When both left and right cell populations express the same CAM (nectin-1), they do not mix or intercalate at the boundary, since the adhesive forces between the two populations are balanced equally, and hence cancel out (Figure 5b). In contrast, when the two populations express different CAMs (nectin-1 or nectin-3), the difference in self-adhesive forces results in the mixing and intercalation pattern at the boundary (Figure 5c). The simulation hence shows how either intercalation or smooth boundaries can arise at the interface of two growing cell populations depending on the expression levels and adhesive properties of their CAMs.

**Figure 5.**
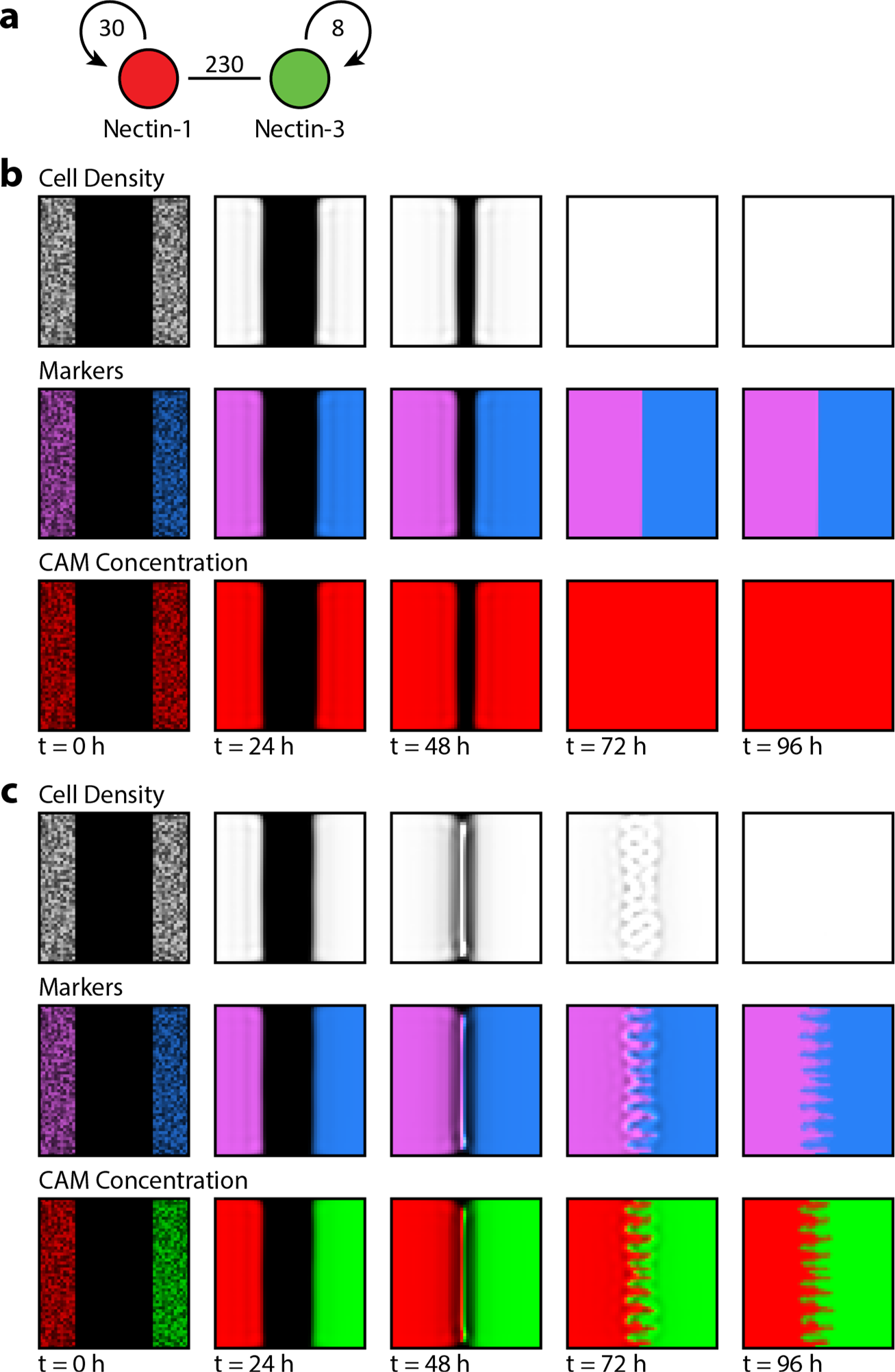
Simulation of proliferating cell populations expressing identical or different CAMs recapitulate *in vitro* intercalation behaviors. The two cell populations express a different marker (magenta or blue), and either the same or different CAMs (red and green). (**a**) Self- and cross-adhesive strengths of nectin-1 and nectin-3. (**b**) Cell populations proliferating and expressing the same CAM, nectin-1, do not mix when they meet at the boundary. (**c**) Cell populations proliferating and expressing different CAMs, nectin-1 (red) or nectin-3 (green), mix and form intercalated patterns when they meet at the boundary due to the different self-adhesion strengths of the two CAMs expressed. Domain of size 1 × 1 mm, discretized into a 50 × 50 grid.

### 3.3. Dynamic regulation of adhesion during gastrulation

During gastrulation, Nodal acts as a diffusive morphogen forming a concentration gradient which induces mesendoderm differentiation (39). In zebrafish, Nodal is expressed in the yolk syncytial layer (YSL), a region of the yolk consisting of nuclei that have descended from the blastoderm (40). The YSL is divided into two segments: the internal YSL (iYSL), which is completely covered by the blastoderm, and the external YSL (eYSL), which protrudes beyond the blastoderm margin. Only the nuclei of the eYSL are transcriptionally active, being the source of the Nodal signal that diffuses through a small area of the blastoderm at the region of the embryonic shield. All germ layers express similar levels of E-cadherin, but Nodal induces the up-regulation in expression of N-cadherin (41). These Nodal-induced cells with higher expression of N-cadherin have higher cell adhesion strengths compared to ectoderm cells (18) and they differentiate into mesendoderm. Those not exposed to the Nodal gradient become ectoderm (42). Furthermore, the Nodal-induced cells with higher N-cadherin levels that differentiate into mesendoderm involute over the blastoderm margin towards the yolk (40). Figure 6 shows the zebrafish gastrulation between the 40% epiboly and shield stages, showing the involution movement of Nodal-induced cells towards the blastoderm margin.

**Figure 6.**
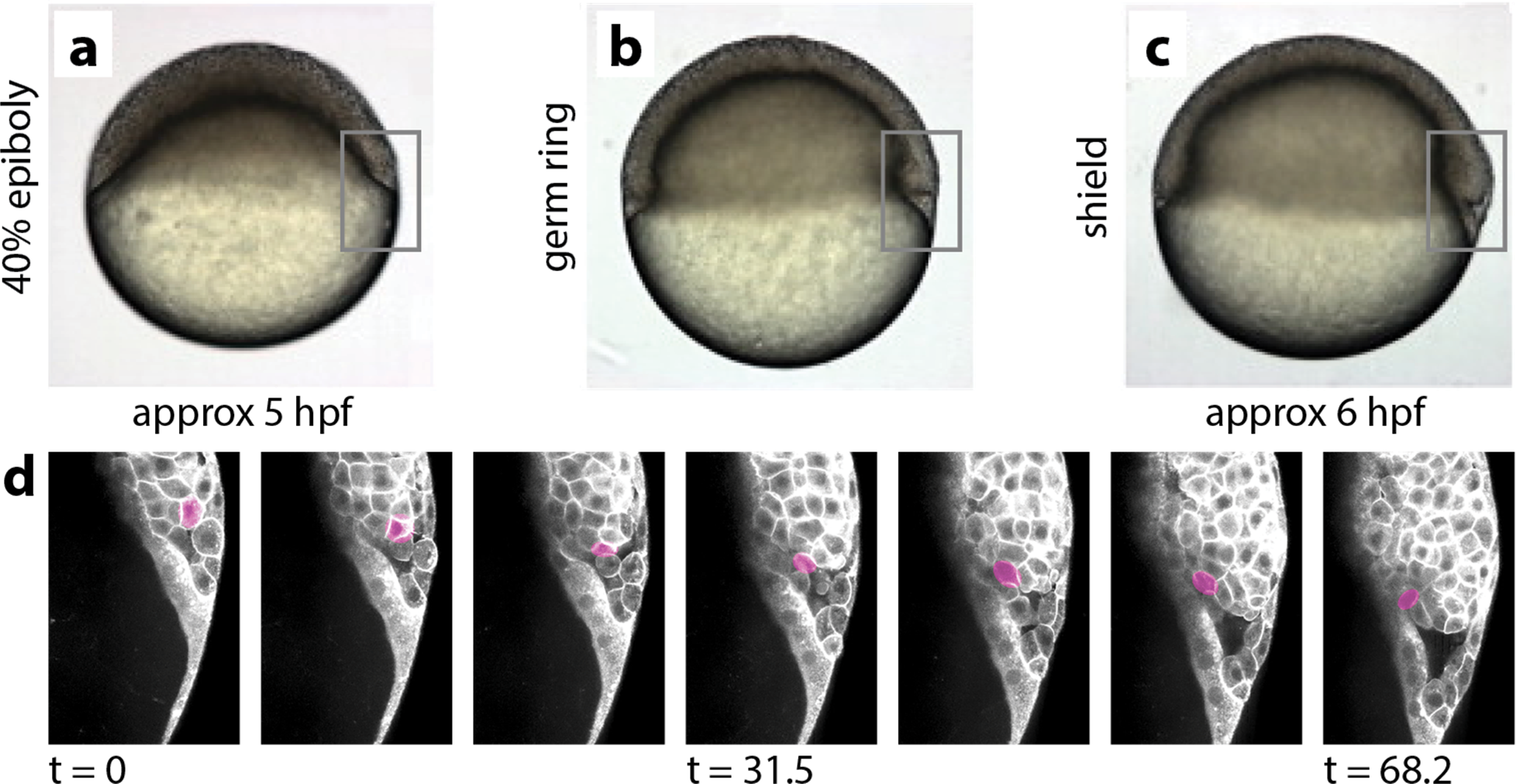
Involution cell movements during zebrafish gastrulation. **(a-c)** Zebrafish embryos during gastrulation at 40% epiboly (a), germ ring (b), and shield (c) stages. **(d)** Sagittal views during gastrulation at the region of the shield from 40% epiboly to shield stages. A manually-tracked single cell (magenta) shows the involution movement of the germ layer cells. Top row adapted from (62), time in hours post fertilization (hpf); bottom row adapted from (63), time in minutes.

We extend the nondimensional model (2.11) to include a regulatory term for the CAMs and equations for the Nodal morphogen and other regulatory factors in the system, such that

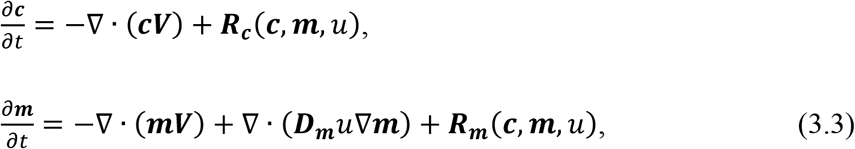

where ***c*** is the vector of CAMs densities, ***R***_***c***_ (***c***, ***m***, *u*) is the regulation of CAMs expression, ***m*** is the vector of morphogen (and other factors) concentrations, ***D***_***m***_ is the vector of diffusion constants per unit of cell density of the morphogens, and ***R***_***m***_ (***c***, ***m***, *u*) is the regulation of morphogens expression.

We employ this model to simulate zebrafish gastrulation due to the dynamic regulation of CAMs by the diffusion of morphogen Nodal (Figure 7). The model includes four CAMs: E-cadherin, N-cadherin, and those present in the EVL and yolk (labeled EVL and Yolk in Figure 7, for simplicity).

**Figure 7.**
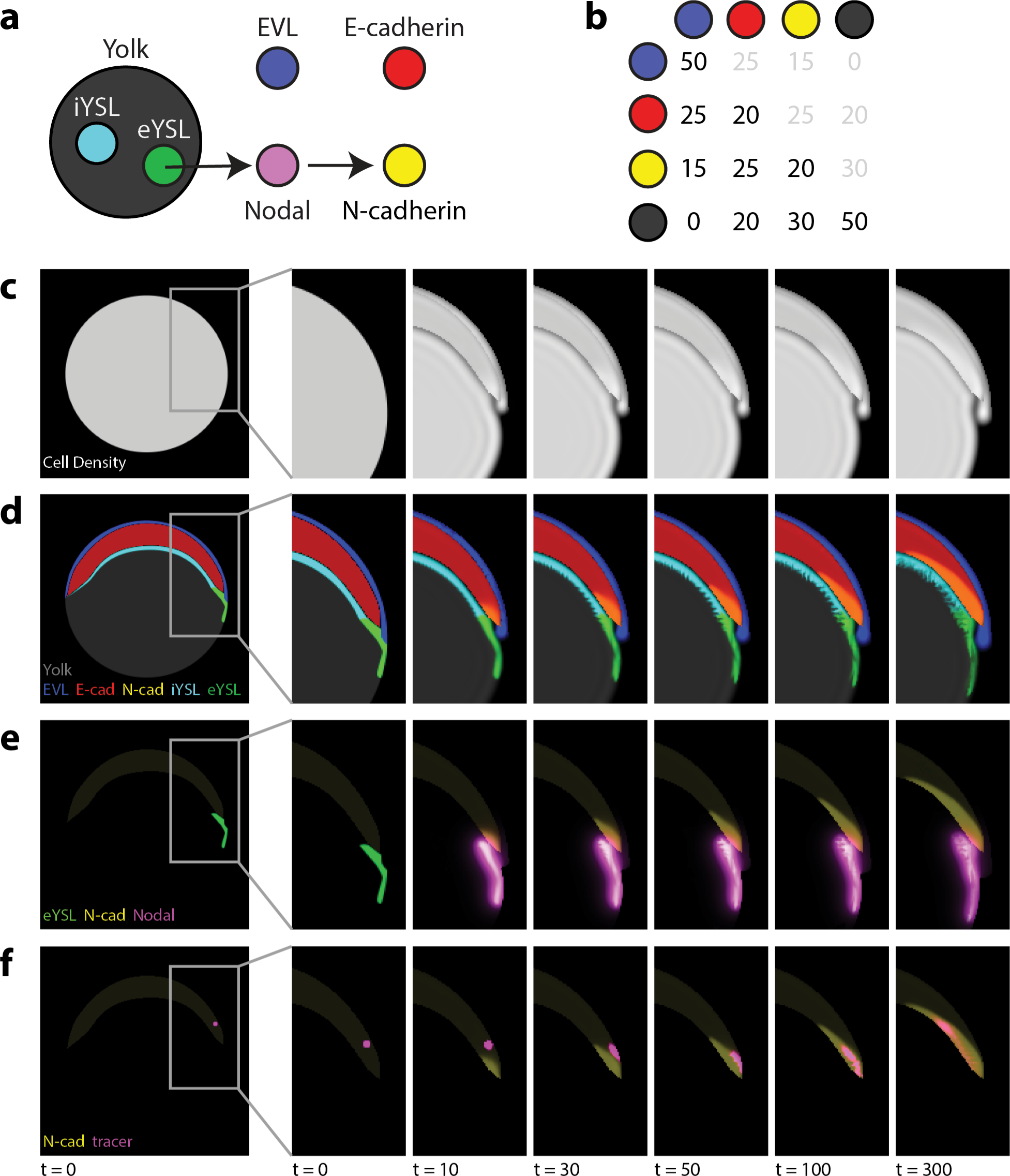
Simulation of zebrafish gastrulation due to the dynamic regulation of CAM expression. **(a)** Regulatory network in the system. iYSL and eYSL regions located in the yolk syncytial layer express non-diffusible factors labeled with the same name (cyan and green, respectively). eYSL factors induce the expression of the diffusible protein Nodal (pink), which induces a higher expression of adhesion protein N-cadherin (yellow). The cells in the enveloping layer (EVL) produce specific CAMs (blue). All germ layers express the same level of E-cadherin adhesion protein (red). **(b)** Adhesion strengths between CAMs. **(c)** The cell density in the embryo is initialized homogenously, but dynamically changes due to differences in cell adhesion. **(d)** Mesendodermal progenitors involute from the germ ring towards the animal pole over the margin of the yolk due to the dynamic up-regulation of N-cadherin expression. **(e)**Nodal expressed due to eYSL factors in the yolk diffuses to the blastoderm, inducing higher levels of N-cadherin expression in this area, which results in the cells moving towards the animal pole beyond the area of Nodal diffusion. **(f)** A non-diffusible tracer (pink) is advected by the mesendodermal progenitor cells, showing their involution. The initial state values are zero except at the locations shown (c-f, *t* = 0), which are homogeneous with *u*_0_ = 0.8, *c*_*yolk*,0_ =0.8, *c*_*evl*,0_ = 0.8, *c*_*Ecad*,0_ = 0.64, *c*_*Ncad*,0_ = 0.16, *m*_*iYSL*,0_ = 0.8 and *m*_*eYSL*,0_ = 0.8, *m*_*nodal*,0_ = 0, and *m*_*tracer*,0_ = 1. Nondimensional parameter values *k*_*r*_ = 5, *D*_*nod*_ = 1, *k*_*e*_ = 10, *λ* = 5. Domain of size 25 × 25 units, discretized into a 250 × 250 grid. Arbitrary units.

The adhesive constants between the CAMs are shown in Figure 7b. E-cadherin and N-cadherin have the same self- and cross-adhesion strengths, but E-cadherin has a stronger adhesion strength with the EVL in comparison to N-cadherin. The CAMs regulatory terms in ***R***_***c***_ (***c***, ***m***, *u*) are all zero, except for N-cadherin, which is regulated by the morphogen Nodal. Hence, the expression of N-cadherin depends on the levels of the morphogen Nodal, in addition to the cell density (more cells express more proteins) and a logistic saturation term, such that

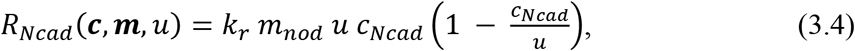

where *k*_*r*_ is a regulatory constant, and *m*_*nod*_ is the concentration of Nodal. In addition to the morphogen Nodal, the model includes the regulatory factors expressed in the iYSL and eYSL regions of the yolk as two lumped variables in ***m*** with zero diffusion constant and regulation. Nodal diffuses and is expressed in the eYSL, hence its diffusion constant is not zero, and its reaction term depends on the eYSL factors in addition to natural degradation, such that

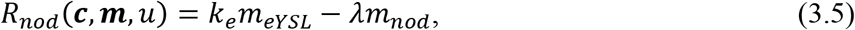

where *k*_*e*_ is an expression constant, *m*_*eYSL*_ is the concentration of eYSL factors, and *λ* is the decay constant for Nodal.

Substituting (3.5) and (3.4) in (3.3), the zebrafish gastrulation model is defined with

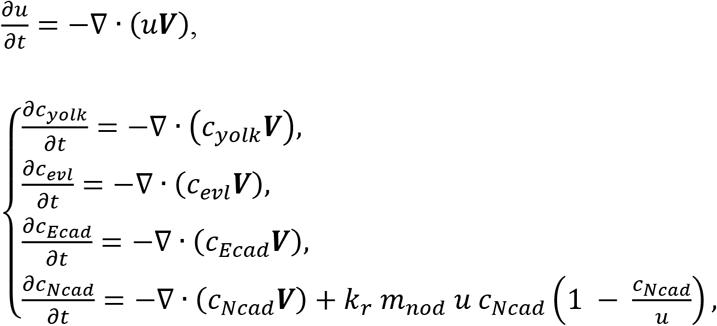

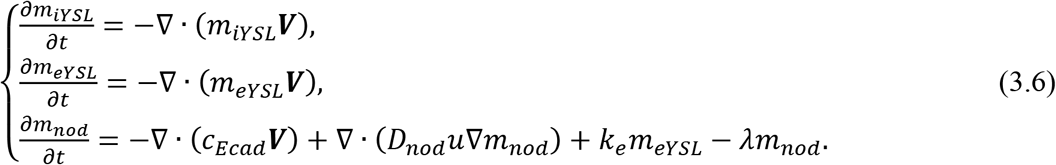

The simulation initial state is based on the experimental images of zebrafish gastrulation in Figure 6a and d. The embryo is defined with a homogeneous cell density circle (Figure 7c, *t* = 0). Within the embryo, the initial EVL and Yolk CAMs densities are set at the locations of the EVL and yolk, respectively, while the E-cadherin and N-cadherin are set in the blastoderm area, with E-cadherin being higher than N-cadherin (Figure 7d, *t* = 0), as it has been shown experimentally (18, 41). The factors iYSL and eYSL are initialized homogeneously in their respective regions (Figure 7d, *t* = 0), while Nodal is initially zero along all the domain (Figure 7e, *t* = 0). A zero-flux boundary was imposed at the interface between the blastoderm with the EVL and yolk regions, to simulate the sealing between EVL cells via apical junctional complexes (43) and the dense cortical yolk cell cytoskeleton (44), respectively. For simplicity, the simulation does not include the EVL cell migration over the yolk.

Figure 7c-f shows the simulation of zebrafish gastrulation, during which the mesendoderm progenitors involute over the margin of the yolk due to the dynamic regulation of CAM expression by the diffusing morphogen Nodal. The eYSL factors activate the expression of Nodal at the eYSL region of the yolk, which then diffuses through the germ ring region of the blastoderm, causing the upregulation of N-cadherin in the cells at this location (Figure 7e). The higher expression of N-cadherin causes a net change of adhesive forces in these cells, which produces a movement towards and upwards the margin of the yolk. This movement extends beyond the diffusion region of Nodal, and results in the involution from the germ ring towards the animal pole. A non-diffusible tracer in the simulation (Figure 7f) shows the involution movement of the cells and their movement along the margin of the yolk, which recapitulates what is seen in the experimental data (Figure 6d). In this way, the model shows how the dynamic regulation of CAMs expression can cause specific patterns and cellular behaviors during gastrulation that are essential for the correct development of the embryo.

## 4. Discussion

This paper presented a novel continuous mathematical model of cell-cell adhesion due to the explicit expression and regulation of cell adhesion proteins (CAMs). Cells express CAMs, which have specific adhesive properties when binding to CAMs of the same type, as well as when binding to CAMs of a different type. The adhesive forces between CAMs produce cell movements dependent on the cell density and CAM concentration of neighbor cells within a sensing radius. Moving cells carry their expressed CAMs and other factors with them. CAM expression can be regulated by intra- and extra-cellular factors, such as diffusible morphogens like Nodal. As regulation alters the expression level of specific CAMs, the adhesive properties of cells are defined dynamically. In this way, the regulatory dynamics of CAM expression can dictate the resulting tissue patterns and shapes.

The capacity of the model to recapitulate cellular *in vitro* and *in vivo* behaviors were demonstrated with three sets of numerical experiments. First, we showed how this approach could recapitulate the classical sorting behaviors in a model of spatially-randomized cells expressing one of two different CAMs. The simulations showed how the emergent sorting dynamics displayed by the cell populations—either engulfment, partial engulfment, mixing, or complete sorting—depended on the cross- and self-adhesion strengths between the expressed CAMs. Next, we showed how intercalation dynamics in a growing cell population depended on the type of CAMs expressed and their adhesion strengths. When the two proliferating populations expressed the same CAM, they formed a completely separated boundary between them. However, when the two proliferating populations expressed different CAMs, they intercalated at the boundary, forming a mixed pattern. Importantly, the regulation of CAMs can be explicitly included in the proposed model, as it is essential in many *in vivo* behaviors. We demonstrated this capacity in the last experiment, which showed cellular involution behaviors controlled by a diffusible morphogen during zebrafish gastrulation. The model included the expression and diffusion of the morphogen Nodal, which formed a concentration gradient extending towards the blastoderm. Nodal induced the up regulation of a specific cadherin, implicitly changing the adhesive properties of these cells. These new adhesive properties resulted in cellular involution over the margin of the yolk, a movement that continued beyond the Nodal gradient due to the properties of the new CAM expression. In summary, these simulations show the ability of the proposed model to explain how dynamic regulation of CAMs can alter cell-cell adhesion properties, producing emergent spatial patterns at the level of cells, tissues, and the whole organism.

Previous continuous modeling approaches have replicated cellular behaviors due to cell-cell adhesion by modeling cell types with specific adhesion properties. Continuous models of cell-cell adhesion have demonstrated cell sorting (26) and intercalation (27) dynamics by explicitly modeling two distinct cellular populations with different cell-cell adhesion constants. Similarly, proposed continuous model of tumor dynamics include static adhesion coefficients defined between cancer cells and the extracellular matrix (29). Other continuous approaches have modeled separated cellular populations representing different cell types, where one cell population could transition to another one as when acquiring a mutation (32), but still the adhesive properties between each population are pre-defined and static. While these models are excellent approaches for simulating large cellular populations with static adhesive properties, they are limited in their capacity to model dynamic adhesive behaviors. In contrast, the continuous model presented here can recapitulate the behaviors of both cell populations with static adhesive properties (simulations 1 and 2) and those with dynamically regulated adhesive properties (simulation 3) by directly including the expression and concentrations of CAMs. This dynamic regulation of CAMs is a key element in many biological processes, and to our knowledge has previously only been captured with hybrid models (25). However, numerically solving hybrid models are computationally infeasible for large cell numbers and their mathematical analysis is limited due to their discrete nature.

The proposed approach uses adhesion forces that arise dynamically from CAM expression, instead from explicit cell types. Single-cell force spectroscopy can directly measure the adhesion forces between cells expressing different levels of CAMs (45), which can be used to experimentally set the parameters of the model. For simplicity, the model assume that the binding forces are proportional to the product of the relative concentration of CAMs; however, more complex formulations of adhesion ligands and receptors are also possible (46). The extracellular matrix is an additional important element in cell-cell adhesion dynamics and it could be incorporated into the continuous model of adhesion (29). Together with adhesion forces, cell cortex elasticity and tension also can play a role in certain behaviors and be essential during tissue shape dynamics (18, 47). Future work will extend the presented model to incorporate the role of these components in cellular behaviors.

The regulation of CAM expression and how these molecules affect large scale cellular behaviors is extraordinary important in both healthy and diseased states (48–50). The proposed model integrates genetic regulation of CAMs, the biophysical forces of adhesion that drive cell motion, and the subsequent cellular dynamics. This integrated view of dynamic adhesion will be essential for understanding tissue behaviors in developmental and cancer biology, as well as in bioengineering (51, 52). The proposed continuum approach allows for the simulation of large cell populations (up to whole-organism scale), as well as the continuous phenomena involved in genetic regulation (e.g. morphogen gradients and CAM expression). Crucially, machine learning approaches for the reverse-engineering of the regulation of patterning (53–55) and cancer formation (56, 57) directly from formalized experimental data (58–61) can be integrated with the proposed model, with the goal to discover the specific regulatory mechanisms of CAMs that give rise to key spatial phenotypes. In summary, the presented continuous modeling approach will pave the way for the understanding of the regulatory dynamics of cell adhesion essential in developmental, regenerative, and cancer biology.

## Author contributions

J.M.K. and D.L. designed the study, developed the models, and wrote the manuscript; J.M.K. performed the simulations; D.L. secured funding.

## Acknowledgments

We thank Bradford Peercy for critical comments on the article and the members of the Lobo Lab for helpful discussions. This work was supported by the National Science Foundation (NSF) under grant IIS-1566077. Computations used the UMBC High Performance Computing Facility (HPCF) supported by the NSF MRI program grants OAC-1726023, CNS-0821258, and CNS-1228778, the SCREMS program grant DMS-0821311, and UMBC.

## References

1. Foty, R.A., and M.S. Steinberg. 2013. Differential adhesion in model systems. Wiley Interdiscip. Rev. Dev. Biol. 2: 631–645.

2. Townes, P.L., and J. Holtfreter. 1955. Directed movements and selective adhesion of embryonic amphibian cells. J. Exp. Zool. 128: 53–120.

3. Steinberg, M.S. 1958. On the Chemical Bonds between Animal Cells. A Mechanism for Type-Specific Association. Am. Nat. 92: 65–81.

4. Maître, J.-L., and C.-P. Heisenberg. 2013. Three Functions of Cadherins in Cell Adhesion. Curr. Biol. 23: R626–R633.

5. Samanta, D., and S.C. Almo. 2015. Nectin family of cell-adhesion molecules: Structural and molecular aspects of function and specificity. Cell. Mol. Life Sci. 72: 645–658.

6. Togashi, H., K. Kominami, M. Waseda, H. Komura, J. Miyoshi, M. Takeichi, and Y. Takai. 2011. Nectins Establish a Checkerboard-Like Cellular Pattern in the Auditory Epithelium. Science. 333: 1144–1147.

7. Bruner, H.C., and P.W.B. Derksen. 2018. Loss of E-Cadherin-Dependent Cell–Cell Adhesion and the Development and Progression of Cancer. Cold Spring Harb. Perspect. Biol. 10: a029330.

8. Fuhrmann, A., A. Banisadr, P. Beri, T.D. Tlsty, and A.J. Engler. 2017. Metastatic State of Cancer Cells May Be Indicated by Adhesion Strength. Biophys. J. 112: 736–745.

9. Thiery, J.P., H. Acloque, R.Y.J. Huang, and M.A. Nieto. 2009. Epithelial-Mesenchymal Transitions in Development and Disease. Cell. 139: 871–890.

10. Xing, J., and X.-J. Tian. 2019. Investigating epithelial-to-mesenchymal transition with integrated computational and experimental approaches. Phys. Biol. 16: 031001.

11. Wada, N. 2011. Spatiotemporal changes in cell adhesiveness during vertebrate limb morphogenesis. Dev. Dyn. 240: 969–978.

12. Ahrens, M.J., Y. Li, H. Jiang, and A.T. Dudley. 2009. Convergent extension movements in growth plate chondrocytes require gpi-anchored cell surface proteins. Development. 136: 3463–3474.

13. Seguin, L., J.S. Desgrosellier, S.M. Weis, and D.A. Cheresh. 2015. Integrins and cancer: Regulators of cancer stemness, metastasis, and drug resistance. Trends Cell Biol. 25: 234–240.

14. Hiratsuka, S., S. Goel, W.S. Kamoun, Y. Maru, D. Fukumura, D.G. Duda, and R.K. Jain. 2011. Endothelial focal adhesion kinase mediates cancer cell homing to discrete regions of the lungs via E-selectin up-regulation. Proc. Natl. Acad. Sci. 108: 3725–3730.

15. Tai, Y., L. Chen, and T. Shen. 2015. Emerging Roles of Focal Adhesion Kinase in Cancer. Biomed Res. Int. 2015: 1–13.

16. Graner, F., and J.A. Glazier. 1992. Simulation of biological cell sorting using a two-dimensional extended Potts model. Phys. Rev. Lett. 69: 2013–2016.

17. Glazier, J.A., and F. Graner. 1993. Simulation of the differential adhesion driven rearrangement of biological cells. Phys. Rev. E. 47: 2128–2154.

18. Krieg, M., Y. Arboleda-Estudillo, P.H. Puech, J. Käfer, F. Graner, D.J. Müller, and C.P. Heisenberg. 2008. Tensile forces govern germ-layer organization in zebrafish. Nat. Cell Biol. 10: 429–436.

19. Zajac, M., G.L. Jones, and J.A. Glazier. 2003. Simulating convergent extension by way of anisotropic differential adhesion. J. Theor. Biol. 222: 247–259.

20. Gerisch, A., and K.J. Painter. 2010. Mathematical modeling of cell adhesion and its applications to developmental biology and cancer invasion. In: Chauvière A, L Preziosi, C Verdier, editors. Cell Mechanics: From Single Scale-Based Models to Multiscale Modelling. CRC Press. pp. 319–350.

21. Glazier, J.A., Y. Zhang, M. Swat, B. Zaitlen, and S. Schnell. 2008. Coordinated Action of N-CAM, N-cadherin, EphA4, and ephrinB2 Translates Genetic Prepatterns into Structure during Somitogenesis in Chick. Curr. Top. Dev. Biol. 81: 205–247.

22. Turner, S., and J.A. Sherratt. 2002. Intercellular adhesion and cancer invasion: A discrete simulation using the extended potts model. J. Theor. Biol. 216: 85–100.

23. Anderson, A.R.A. 2005. A hybrid mathematical model of solid tumour invasion: the importance of cell adhesion. Math. Med. Biol. 22: 163–186.

24. Zhang, Y., G.L. Thomas, M. Swat, A. Shirinifard, and J.A. Glazier. 2011. Computer simulations of cell sorting due to differential adhesion. PLoS One. 6: 26–28.

25. Ramis-Conde, I., D. Drasdo, A.R.A. Anderson, and M.A.J. Chaplain. 2008. Modeling the influence of the E-cadherin-β-catenin pathway in cancer cell invasion: A multiscale approach. Biophys. J. 95: 155–165.

26. Armstrong, N.J., K.J. Painter, and J.A. Sherratt. 2006. A continuum approach to modelling cell-cell adhesion. J. Theor. Biol. 243: 98–113.

27. Murakawa, H., and H. Togashi. 2015. Continuous models for cell–cell adhesion. J. Theor. Biol. 374: 1–12.

28. Armstrong, N.J., K.J. Painter, and J.A. Sherratt. 2009. Adding adhesion to a chemical signaling model for somite formation. Bull. Math. Biol. 71: 1–24.

29. Gerisch, A., and M.A.J. Chaplain. 2008. Mathematical modelling of cancer cell invasion of tissue: Local and non-local models and the effect of adhesion. J. Theor. Biol. 250: 684–704.

30. Kim, Y., S. Lawler, M.O. Nowicki, E.A. Chiocca, and A. Friedman. 2009. A mathematical model for pattern formation of glioma cells outside the tumor spheroid core. J. Theor. Biol. 260: 359–371.

31. Painter, K.J., N.J. Armstrong, and J.A. Sherratt. 2010. The impact of adhesion on cellular invasion processes in cancer and development. J. Theor. Biol. 264: 1057–1067.

32. Domschke, P., D. Trucu, A. Gerisch, and M.A.J. Chaplain. 2014. Mathematical modelling of cancer invasion: Implications of cell adhesion variability for tumour infiltrative growth patterns. J. Theor. Biol. 361: 41–60.

33. Gerisch, A. 2010. On the approximation and efficient evaluation of integral terms in PDE models of cell adhesion. IMA J. Numer. Anal. 30: 173–194.

34. Weiner, R., B.A. Schmitt, and H. Podhaisky. 1997. ROWMAP--a ROW-code with Krylov techniques for large stiff ODEs. Appl. Numer. Math. 25: 303–319.

35. Steinberg, M.S. 1963. Reconstruction of Tissues by Dissociated Cells. Science. 141: 401–408.

36. Cochet-Escartin, O., T.T. Locke, W.H. Shi, R.E. Steele, and E.-M.S. Collins. 2017. Physical Mechanisms Driving Cell Sorting in Hydra. Biophys. J. 113: 2827–2841.

37. Armstrong, P.B. 1971. Light and electron microscope studies of cell sorting in combinations of chick embryo neural retina and retinal pigment epithelium. Wilhelm Roux. Arch. Entwickl. Mech. Org. 168: 125–141.

38. Harrison, O.J., J. Vendome, J. Brasch, X. Jin, S. Hong, P.S. Katsamba, G. Ahlsen, R.B. Troyanovsky, S.M. Troyanovsky, B. Honig, and L. Shapiro. 2012. Nectin ectodomain structures reveal a canonical adhesive interface. Nat. Struct. Mol. Biol. 19: 906–915.

39. Schier, A.F. 2009. Nodal Morphogens. Cold Spring Harb. Perspect. Biol. 1: a003459–a003459.

40. Carvalho, L., and C.-P. Heisenberg. 2010. The yolk syncytial layer in early zebrafish development. Trends Cell Biol. 20: 586–592.

41. Giger, F.A., and N.B. David. 2017. Endodermal germ-layer formation through active actin-driven migration triggered by N-cadherin. Proc. Natl. Acad. Sci. 114: 201708116.

42. Rodaway, a, H. Takeda, S. Koshida, J. Broadbent, B. Price, J.C. Smith, R. Patient, and N. Holder. 1999. Induction of the mesendoderm in the zebrafish germ ring by yolk cell-derived TGF-beta family signals and discrimination of mesoderm and endoderm by FGF. Development. 126: 3067–3078.

43. Stemmler, M.P., B. Koschorz, T.J. Carney, H. Schwarz, A. Amsterdam, M. Hammerschmidt, and K. Slanchev. 2009. The Epithelial Cell Adhesion Molecule EpCAM Is Required for Epithelial Morphogenesis and Integrity during Zebrafish Epiboly and Skin Development. PLoS Genet. 5: e1000563.

44. Bruce, A.E.E. 2016. Zebrafish epiboly: Spreading thin over the yolk. Dev. Dyn. 245: 244–258.

45. Malek-Zietek, K.E., M. Targosz-Korecka, and M. Szymonski. 2017. The impact of hyperglycemia on adhesion between endothelial and cancer cells revealed by single-cell force spectroscopy. J. Mol. Recognit. 30.

46. Xu, H., and D.E. Shaw. 2016. A Simple Model of Multivalent Adhesion and Its Application to Influenza Infection. Biophys. J. 110: 218–233.

47. Manning, M.L., R.A. Foty, M.S. Steinberg, and E.-M. Schoetz. 2010. Coaction of intercellular adhesion and cortical tension specifies tissue surface tension. Proc. Natl. Acad. Sci. 107: 12517–12522.

48. Iwamoto, D. V., and D.A. Calderwood. 2015. Regulation of integrin-mediated adhesions. Curr. Opin. Cell Biol. 36: 41–47.

49. Teo, W.W., V.F. Merino, S. Cho, P. Korangath, X. Liang, R.C. Wu, N.M. Neumann, A.J. Ewald, and S. Sukumar. 2016. HOXA5 determines cell fate transition and impedes tumor initiation and progression in breast cancer through regulation of E-cadherin and CD24. Oncogene. 35: 5539–5551.

50. Ito, K., S.H. Park, A. Nayak, J.H. Byerly, and H.Y. Irie. 2016. PTK6 inhibition suppresses metastases of triple-negative breast cancer via SNAIL-dependent E-cadherin regulation. Cancer Res. 76: 4406–4417.

51. Glass, D.S., and I.H. Riedel-Kruse. 2018. A Synthetic Bacterial Cell-Cell Adhesion Toolbox for Programming Multicellular Morphologies and Patterns. Cell. 174: 649–658.e16.

52. Toda, S., L.R. Blauch, S.K.Y.Y. Tang, L. Morsut, W.A. Lim, S. Biology, S. Cell, and L. Angeles. 2018. Programming self-organizing multicellular structures with synthetic cell-cell signaling. Science. 0271: 1–13.

53. Lobo, D., and M. Levin. 2015. Inferring regulatory networks from experimental morphological phenotypes: a computational method reverse-engineers planarian regeneration. PLoS Comput. Biol. 11: e1004295.

54. Lobo, D., J. Morokuma, and M. Levin. 2016. Computational discovery and in vivo validation of hnf4 as a regulatory gene in planarian regeneration. Bioinformatics. 32: 2681–2685.

55. Lobo, D., and M. Levin. 2017. Computing a Worm: Reverse-Engineering Planarian Regeneration. In: Adamatzky A, editor. Advances in Unconventional Computing. Volume 2: Prototypes, Models and Algorithms. Springer International Publishing. pp. 637–654.

56. Lobikin, M., D. Lobo, D.J. Blackiston, C.J. Martyniuk, E. Tkachenko, and M. Levin. 2015. Serotonergic regulation of melanocyte conversion: A bioelectrically regulated network for stochastic all-or-none hyperpigmentation. Sci. Signal. 8: ra99.

57. Lobo, D., M. Lobikin, and M. Levin. 2017. Discovering novel phenotypes with automatically inferred dynamic models: a partial melanocyte conversion in Xenopus. Sci. Rep. 7: 41339.

58. Lobo, D., E.B. Feldman, M. Shah, T.J. Malone, and M. Levin. 2014. A bioinformatics expert system linking functional data to anatomical outcomes in limb regeneration. Regeneration. 1: 37–56.

59. Lobo, D., T.J. Malone, and M. Levin. 2013. Towards a bioinformatics of patterning: a computational approach to understanding regulative morphogenesis. Biol. Open. 2: 156–169.

60. Lobo, D., T.J. Malone, and M. Levin. 2013. Planform: an application and database of graph-encoded planarian regenerative experiments. Bioinformatics. 29: 1098–1100.

61. Lobo, D., E.B. Feldman, M. Shah, T.J. Malone, and M. Levin. 2014. Limbform: a functional ontology-based database of limb regeneration experiments. Bioinformatics. 30: 3598–3600.

62. Lachnit, M., E. Kur, and W. Driever. 2008. Alterations of the cytoskeleton in all three embryonic lineages contribute to the epiboly defect of Pou5f1 / Oct4 deficient MZ spg zebrafish embryos. Dev. Biol. 315: 1–17.

63. Montero, J.-A.J.-A., L. Carvalho, M. Wilsch-Bräuninger, B. Kilian, C. Mustafa, and C.-P. Heisenberg. 2005. Shield formation at the onset of zebrafish gastrulation. Development. 132: 1187–1198.

